# Native Triboelectric Nanogenerator Ion Mobility-Mass Spectrometry of Egg Proteins Relevant to Objects of Cultural Heritage at Picoliter and Nanomolar Quantities

**DOI:** 10.1101/2023.01.09.523258

**Authors:** Daniel D. Vallejo, Aleksandra Popowich, Julie Arslanoglu, Caroline Tokarski, Facundo M. Fernández

## Abstract

Native mass spectrometry (nMS) has found widespread success in measuring native-like protein structures in the gas-phase and, when combined with ion mobility (IM), is capable of measuring protein collision cross sections (CCS) and stabilities. These methods are well validated, but often rely on samples that are abundantly available through repeated recombinant protein expression. For ultra-precious and irreplaceable samples from cultural heritage objects, protein content can be far below the micromolar and microliter levels required for robust protein experiments, a major hurdle in characterizing protein higher order structure and degradation mechanisms. Combining triboelectric nanogenerators (TENG) and IM-MS enables measuring protein size and stability rapidly from ultra-small sample quantities. Here, TENG IM-MS is implemented with standard and sub-micron emitters to characterize proteins relevant to cultural heritage objects, and demonstrates native structures can be obtained even at nanomolar concentrations and picoliter quantities.

## Introduction

Materials in objects of cultural heritage (CH), such as paintings, span several molecular classes ranging from small molecules (mineral pigments and organic dyes) to polymers (proteins, polysaccharides, lipids, synthetics) used as binders, coatings, and adhesives. Characterizing these materials is essential to not only better understand and interpret the artist or maker, but to provide information that may serve to further identify manufacturing and artists’ techniques, clarify attributions, establish sourcing materials, and better preserve the work for future generations. Studying paint binding media and adhesives often involves studying proteins, which have been an important molecular class in objects of CH since the inception of art. These proteins, having numerous sources (e.g. egg, milk, blood, bone, skin, etc.),^1^ have substantial effects on the appearance and stability of paints on canvas and paper paintings, and polychrome sculpture^2^. Despite recent advances in the understanding of proteins found in CH objects^3^ their higher order structure (HOS) remains significantly understudied. HOS is directly correlated to the stability and function of proteins in objects and plays a pivotal role in the object’s integrity and preservation, but a critical hurdle to analysis is the ultra-precious nature of the objects, as well as the low abundance of proteins relative to the total sample.

The technologies used to elucidate the structure of materials in these objects are categorized as either invasive or non-invasive. Non-invasive techniques such as portable Fourier transform infrared (FTIR)^4^ and Raman spectroscopy^5^ are highly attractive because of their ability to extract molecular information without consuming the object’s precious materials. FTIR and Raman are well validated and remain gold-standards for pigment identification, and may be used to determine the localized presence/absence of proteinaceous materials on an objects’ surface. However, the interrogation of protein structures through these techniques remains limited due to the competing signals from other materials in the sample, as well as differential effects that degradation of structure, pigment-interactions, and intrinsic protein fluorophores have on generating reproducible spectroscopic features and often cannot discriminate or identify proteins^6–8^.

When identification of the specific protein must be known, micro-invasive but structurally-sensitive techniques such as mass spectrometry (MS) are employed^3,9^. Most current MS workflows for the analysis of proteins use bottom-up methods^10^ where the protein is enzymatically digested and the peptide sequences analyzed, and likely require front-end separation methods such as chromatography to separate the digested components^11^. While these traditional proteomic techniques are becoming more commonplace in CH as a way to perform protein type, species, or tissue identification, they can be challenging to apply because of their high sample requirements relative to the amount of material that can (or should) be removed from an artwork^3,11^. As such, there is a continued need to develop new MS tools that minimize the consumption of precious samples while providing robust separation from the underlying matrices found in CH objects to provide protein structural information. Along these lines, an ongoing trend in heritage science is to leverage advances in state-of-the-art structural biology technologies, such as foot printing MS techniques (e.g. HDX, cross-linking, etc.)^12^, topdown proteomics, and native mass spectrometry (nMS) to achieve more in-depth protein characterization.

In particular, nMS has become an increasingly valuable tool for structural biology. It has earned this distinction because, unlike bottom-up or top-down proteomic strategies, it can preserve non-covalent interactions of proteins and their complexes. Retaining three-dimensional (tertiary) protein structures and (quaternary) protein-complex structures has enabled characterizing protein ensembles, assemblies, ligand-protein, and protein-protein interaction networks, together with changes in protein stability within such networks. Understandably, many current nMS applications focus on structural biology and disease-related research and, as a result, have greatly benefited from the abundance of readily obtainable protein standards or protein systems that can be recombinantly expressed and purified at yields suitable for routine nMS analysis. However, the amount of sample that can be obtained for MS experiments from a CH object (ideally <10 μg with <10% w/w being protein content) often falls far below the concentration of typical chemical standards (∼10-20 μM) routinely used for robust nMS experiments. Applying these state-of-the-art technologies to CH could address questions regarding the extent to which a protein’s structure changes in objects of cultural heritage.

Triboelectric nanogenerator (TENG)-based electrospray ionization (ESI) MS has recently demonstrated excellent analytical figures of merit. This includes increased sensitivity due to higher electrical fields, reproducibility across a wide range of molecular classes^13,14^, and the ability for synchronous data acquisition with time dispersive instruments for sample-limited analytes^15^. TENG MS has also been used to generate protein ions from model systems^16^, but questions remain regarding whether the protein structures measured are still in their native conformation or if gas-phase structural changes could occur during ion generation.

Here, a large area sliding freestanding inductive TENG ion source hyphenated to an ion mobility (IM)-MS platform is tested for native protein analysis relevant to the CH research field. IM is a technique that separates the gas-phase structures of molecules based on their orientationally-averaged collision cross-section (CCS) values, useful for separating biomolecules from underlying matrices, and has emerged as one of the premier analytical technologies for characterizing the HOS structure, stoichiometry, and biophysical stability of biological macromolecules^17,18^. We demonstrate that TENG IM-MS is capable of generating native protein ions for three protein standards, falling within 3% of their corresponding CCS database values^19^. We also show similar nMS results for eight standard proteins across a molecular weight range of 5 – 800 kDa. Theoretical CCS values were empirically calculated for three egg proteins that have use in CH. Experimental results, with notable exception for lysozyme, were within 3% of their respective theoretical CCS value. Lastly, we demonstrate the analytical capabilities of TENG IM-MS by measuring native ovalbumin at picoliter and nanomolar concentration using both standard and submicron nESI emitters, which is a step forward not only in terms of the quantities of sample needed for standard nMS experiments, but also in making nMS available to the study of objects of CH.

## EXPERIMENTAL SECTION

### Sample Preparation

Protein samples were purchased from Sigma-Aldrich (St. Louis, MO) in lyophilized form. Human Insulin (INS), cytochrome C (CytC), β-lactoglobulin ((β-lac), Bovine Serum Albumin (BSA), Concanavalin A (ConA), Alcohol Dehydrogenase (ADH), Glutamate Dehydrogenase (GDH), and GroEL were used as protein standards for CCS calibration purposes. GroEL was additionally prepared following a previously published protocol^20^. Human Serum Albumin (HSA) was used as a positive control, Albumin from chicken egg (Ovalbumin/OVA), Conalbumin (Ovotransferrin/OVT), and Lysozyme (LYS) from chicken egg white were used as experimental proteins.

Proteins samples were kept on ice and buffer exchanged into 200 mM ammonium acetate buffer at pH 7 using Micro Bio-spin columns with either a 6 or 30 kDa cutoff (Bio-Rad, Hercules, CA). Buffer exchanged samples were then diluted to a working concentration of 10 μM before analysis. A concentration gradient of OVA was made through serial dilution of the 10 μM stock sample to the following concentrations: 7.5 μM, 5 μM, 2.5 μM, 1 μM, 750 nM, 500 nM, 250 nM, and 100 nM.

### Nano-electrospray Ionization (nESl) Conditions

Emitters for nESI experiments were produced in-house from borosilicate thin-wall glass capillaries with an outer diameter (OD) of 1 mm and inner diameter (ID) of 0.78 mm (Harvard Apparatus, Holliston, MA). Borosilicate capillary emitters were prepared with an ID of ∼5 μm using a Sutter Instruments P-97 Flaming Brown Puller (Novato, CA). Additionally, ∼1 μm and submicron needles were pulled using the same instrumentation, and sizes were confirmed using a scanning electron microscope (SEM). Puller programs are provided in the Supplementary Information section (Table S2). Protein samples were loaded into the nESI emitter (2-8 μL), and the emitters were mounted on an x, y, z manual linear stage (Thorlabs, Newton, NJ) to control their position relative to the sampling cone. The emitter tip was held between 7−10 mm away from the inlet orifice. Protein ions were generated in positive ion mode using either a direct current (DC) power supply or a TENG ion source as described below. A Stanford Research Systems (Sunnyvale, CA) High Voltage Power Supply (model PS350/5000V-25W) was used as the DC power supply, operated at voltages of 1.0 - 1.2 kV and a current of 3 μA.

The TENG source ion used was a mechanically-actuated SF-TENGi configuration as described previously^15,16^. Briefly, the static electrode was fabricated with a Cu film deposited onto PTFE as two rectangular regions separated by an uncoated rectangular region on an acrylic support of 135 and 121 cm^2^, and a movable sliding electrode fabricated with Nylon mounted onto a foam square on an acrylic support of 127 cm^2^. A linear motor (LinMot USA, Inc.) was used to operate the movable sliding electrode at a frequency of 1.2 seconds and 15 ms rest time. The copper electrode used in previous TENGi configurations was replaced with an ESI conductive sleeve (Waters, Milford, MA).

### Ion Mobility-Mass Spectrometry

Protein samples were primarily analyzed on a q-traveling wave (TW) IM-TOF Synapt G2 HDMS (Waters, Milford, MA). The sampling cone was operated at 40 V. The backing pressure was set between 2.3-8.1 mbar. The helium cell flow was operated at 200 mL/min and pressurized to 1.40×l0^3^ mbar. The trap traveling-wave ion guide was pressurized to 2.72×10^−2^ mbar of argon gas. The travelling-wave IM separator was operated at a pressure of ∼3.8 mbar. IM separation was achieved with a travelling wave operated at 40 V wave height traveling at 600 m/s. The ToF-MS was operated over the *m/z* range of 1,000–10,000 at a pressure of 1.22×10^−6^ mbar. Additional spectrometer settings are provided in the Supplemental Information (Table S3).

### Collision Induced Unfolding

Protein ions were subjected to collisions in the travelling-wave ion trap prior to the IM separation to perform charge multiplexed CIU. The collision voltage was ramped from 5 to 200 V in 5 V increments to construct CIU fingerprints. The dwell time for each 5 V step was 12 seconds. The IM data was calibrated using the standard reference proteins and ^TW^CCS_N2_ was calculated using the IM-SCal software^21^.

### Data Analysis

Experimental CCS values for the standard proteins were compared to the standard protein CCS Database^19^. Experimental CCS values for HSA, OVA, OVT, and LYS were compared to theoretical CCS values empirically derived from the standard protein CCS database. Additional theoretical CCS values were empirically derived from correlation curves of charge (*z*) vs. CCS or molecular weight (kDa) vs. CCS from the experimental CCS values of the standard proteins. Percent differences were calculated between the CCS database values, and the average CCS experimental values computed from intra- and inter-day CCS measurements. Root-square deviation values were computed for the intra- and inter-day CCS measurements.

Drift time and CCS data were extracted at each collision voltage in Drifts cope (Waters, Milford, MA) using TWIMExtract^22^. These extracted drift time data were analyzed using a home-built software package, ClUSuite 2^23^. The monomeric fingerprints were 2-D smoothed with a Savitzky-Golay function with a smoothing window size of 5 and 2 smooth iterations. The collision voltage axis was interpolated with an axis scaling factor of 2. Standard feature detection was used with a minimum feature length of 3 (steps), an allowed width of 2-2.5 (nm^2^ axis units), and a maximum CV gap length of 0. Ground state and activated arrival time distribution plots were generated by taking the average of three replicates at CVS (ground state) or CV100 (activated) and all intensities were normalized.

## RESULTSAND DISCUSSION

### Method Validation for Native TENG IM-MS

Figure 1A shows the updated design for TENG devices used for ESI and its application towards nMS. The primary modifications included changing the layered materials of the top electrode, as well as the conductive copper tape used to induce charging within the nESI emitter to a commercial carbon sleeve. These changes improved the general reproducibility of the TENG methodology when applied to proteins. Additional details on the nature of these changes as well as their advantages are described in the Methods and Supporting Information sections.

**Figure 1.**
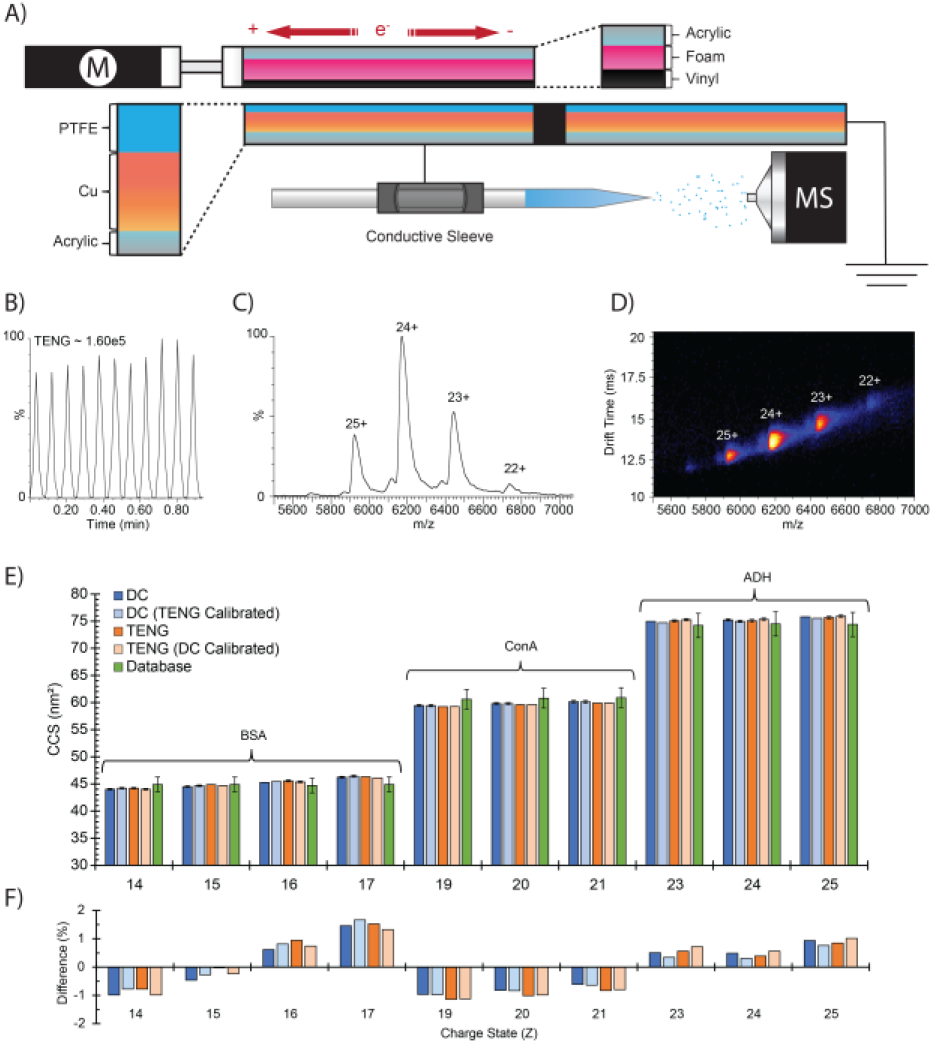
nESI was achieved by alternating between TENG and DC power supplies. (A) the sliding free standing TENG and DC power source were connected to the same needle with the same protein sample. TENG produced comparable (B) total ion currents, (C) mass spectrum and charge state distribution, (D) and IM-MS drift time plots for alcohol dehydrogenase. (E) Additionally, TENG produced comparable CCS calibrated measurements for three standard proteins. These measurements were within ±2% of their corresponding CCS database values.

To evaluate the performance of this newly improved TENG design for nMS, we compared the mass spectral data generated to those generated with a standard DC power supply for three model proteins: BSA, ConA, and ADH. The total ion chronogram, mass spectrum, and drift time plot for ADH are shown in Figures 1B-D. TENG produced nearly identical information content compared to a DC power supply paired with the same nESI emitter (Figure S1). However, as previously discussed in the introduction, charge states alone are not a sufficient indicator for the retention of a proteins native structure since smaller structural alterations (e.g. unfolding) can still be taking place within the same charge states. Using IM, we converted our arrival time distributions to CCS measurements for the accessible charge states for all three model proteins tested (Figure 1E). We obtained the following experimentally averaged CCS measurements: 45.0 ±0.83 nm^2^ for BSA, 59.8 ±.29 nm^2^ for ConA, and 75.33 ±0.35 nm^2^ for ADH with a DC power supply; and 45.2 ±.77 nm^2^ for BSA, 59.6 ±.27 nm^2^ for ConA, and 75.3 ±0.29 nm^2^ for ADH with TENG. To rule out any systematic issues with CCS calibration between the two ionization approaches, we performed a cross calibration test using DC calibrants for TENG data and vice versa; no systematic deviation was observed, and any shifts in calculated CCS values were within error of the normal calibration procedure. Lastly, we compared our experimental results for these three protein standards to a protein CCS database^19^, finding the results within 2% of their corresponding native CCS values (Figure 1F). These results demonstrated that nMS experiments can be conducted using TENG devices without any detectable changes in proteins higher order structures.

### TENG IM-MS of Proteins Relevant to Cultural Heritage

The proteinaceous materials used for cultural heritage objects have numerous sources. This pilot study is based on three proteins sourced from eggs that would be used in art. Eggs have several significant uses in cultural heritage objects as: binders, glues, and varnishes^24,25^; and the proteins in the egg whites having a particular function as varnishes (glair) and binders. Other structural studies that have focused on egg white proteins have demonstrated changes in protein-pigment interactions can be monitored and affect dried paint films, but focused primarily on LYS based systems^12^. The TENG IM-MS system was tested to study three egg-white proteins relevant to cultural heritage: LYS, OVA, and OVT (Figure 2). LYS presented the most complex native mass spectrum, with multiple overlapping oligomeric states (Figure 2A). However, incorporating a drift time dimension simplified the qualitative deconvolution of the monomer (14.4 kDa, 5-7^+^), dimer (9-ll^+^), and trimer oligomeric states for LYS. OVA was mostly present as a monomer (43 kDa, 11-13^+^), with a dimer form in much lower abundance (16-18^+^, ∼5-8%) (Figure 2B). Lastly, OVT predominantly presented as a monomeric form (76 kDa), with a relatively lower abundance dimer (<2%) (Figure 2C). These initial results indicated that TENG IM-MS holds potential towards the study of proteins relevant to CH objects and paintings.

**Figure 2.**
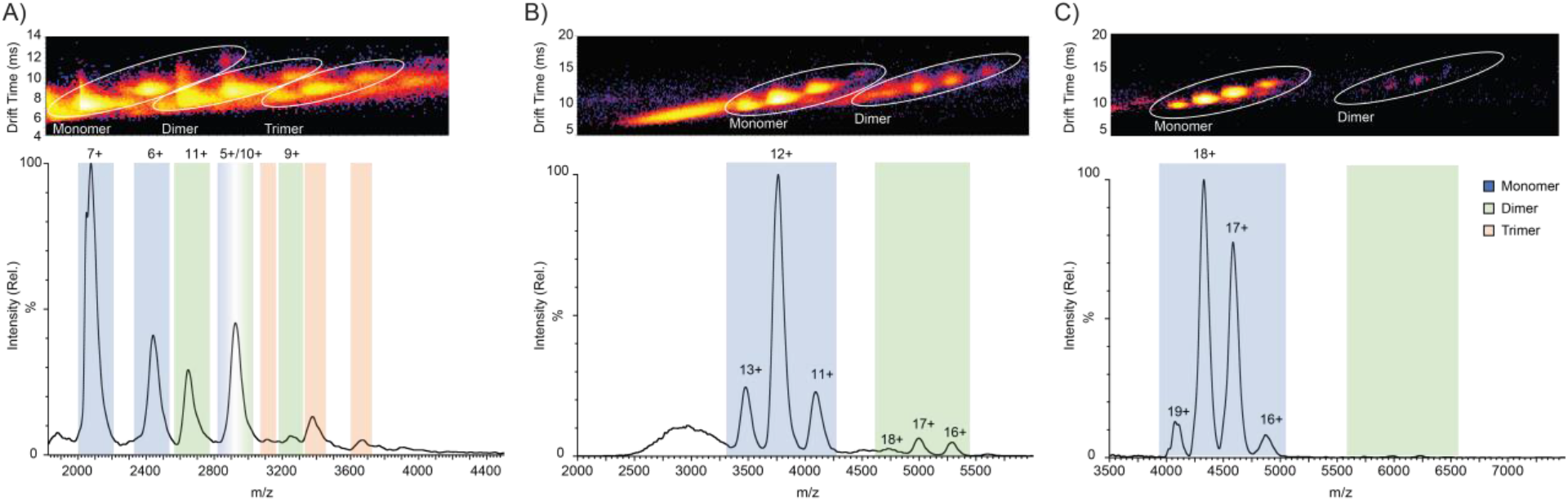
Typical native mass spectra for cultural heritage-relevant proteins from chicken egg white: (A) lysozyme, (B) ovalbumin, (C) and ovotransferrin. Mass spectra and their corresponding IM-MS drift time plots show the distribution of monomer (blue) and dimer (green) species. The trimer was observed for lysozyme (orange).

Following these experiments, we conducted extensive CCS calibration to determine if native structures could be obtained for the egg proteins studied by TENG IM-MS. We expanded the dataset to now include eight standard proteins as described in the methods section. All these proteins have CCS database values available. Comparison of experimental CCS measurements to those in the CCS database showed good agreement (<3.5%) for most of the protein standards except for INS, which had a percent difference of more than ±7% (Table 1). However, there is no CCS value for nitrogen measurements in the CCS database, and the value we used to compare to was based on a helium-to-nitrogen conversion factor that could possibly explain the observed differences (Figure S2). When we instead used an empirically derived CCS value for INS obtained by generating a calibration curve from our CCS measurements of the standard proteins, we obtained better agreement within 3% with our experimental data (Figure 3B).

**Figure 3.**
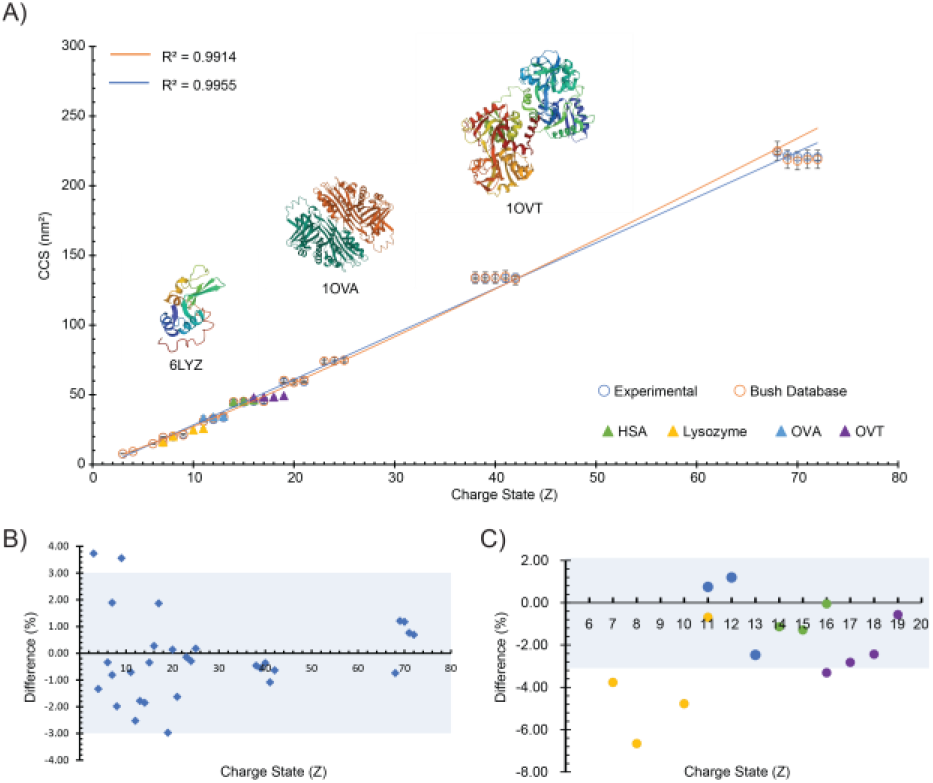
(A) CCS values for eight protein standards from the Bush CCS database (orange curve) and the corresponding experimental CCS measurements for the same standard proteins (blue curve). Correlation plots of charge (z) or molecular weight (kDa) were used to empirically calculated the theoretical CCS of HSA, LYS, OVA, and OVT. (B) Experimental CCS measurements of protein standards compared to their Bush database values showing <3.5% variation. (C) Experimental CCS measurements of egg proteins compared to their theoretical CCS values.

Lastly, we computed theoretical CCS values for the egg proteins using this empirical calibration approach, finding that the theoretical CCS values for the most abundant charge state of LYS, OVA; and OVT to be 166, 336, and 496 nm^2^, respectively. The experimental values obtained were 160 ± 0.8, 340 ± 7.2, and 484 ± 37.2 nm^2^ corresponding to percent differences of-3.77,1.20, and -2.43%, respectively (Figure 3C). Interestingly, larger errors were noted for LYS for all charge states observed. However, LYS has relatively less HOS compared to OVA or OVT, as shown by the crystal structure from the Protein Data Bank (PDB), also embedded in Figure 3A. Initially we believed this structural difference could have resulted in compaction of LYS as it is transferred to the gas-phase, resulting in the larger negative CCS differentials compared to the expected CCS values for a protein of such size or charge. However, chicken egg white LYS CCS values have been reported to range between 13 – 30 nm^2^ under both native and disulfide reducing conditions, indicating it’s wide conformational ensemble^26^’^27^. Additionally, these values are all larger than the calculated cross section for the crystal structure coordinates of Lysozyme (11.8 nm^2^), indicating the diffuse nature of its HOS^27^. Together, this approach provided a path for determining the degree of native structure conserved under TENG IM-nMS conditions.

### Minimal Sample Size

Confirming nMS results for egg proteins is a crucial step towards fully characterizing their structures for the subsequent analysis of similar proteins from objects of CH. However, these measurements used protein standards that are not only sufficiently pure, but also available at volumes exceeding 1 mL and concentrations of ∼10 μM that are not likely obtainable from CH objects. We therefore leveraged the capabilities of sub-nanoliter volume analysis per TENG pulse^15^, and sought to measure the average volume consumption in experiments with model proteins relevant to CH. A single ESI emitter was loaded with 3 μL of OVA at ∼ 10 μM to perform a sample exhaustion TENG nESI experiment. We allowed this experiment to continue uninterrupted until either the sample was completely consumed, or the needle clogged from protein aggregation. It was observed that even after ∼519 minutes of constant TENG IM-MS operation (Figure 4A) the needle never clogged, reaching total sample consumption, as confirmed optically with a microscope. The calculated average number of pulses-per-minute of the experiment was 14, with an average TENG pulse volume of approximately 412 pL, for a total consumed volume of 3 μL. To confirm these results, a duplicate experiment with a different total sample volume (2 μL) was conducted, finding that the average pulse volume was lower, averaging 330 pL (Figure S3B), but similarly the needle did not clog as is often expected in nMS experiments. We hypothesize that the pulsed nature of TENG ESI may produce a sufficiently low flow rate during each pulse that it lowers the propensity of protein aggregation at the tip of the emitter that would typically lead to needle clogging.

**Figure 4.**
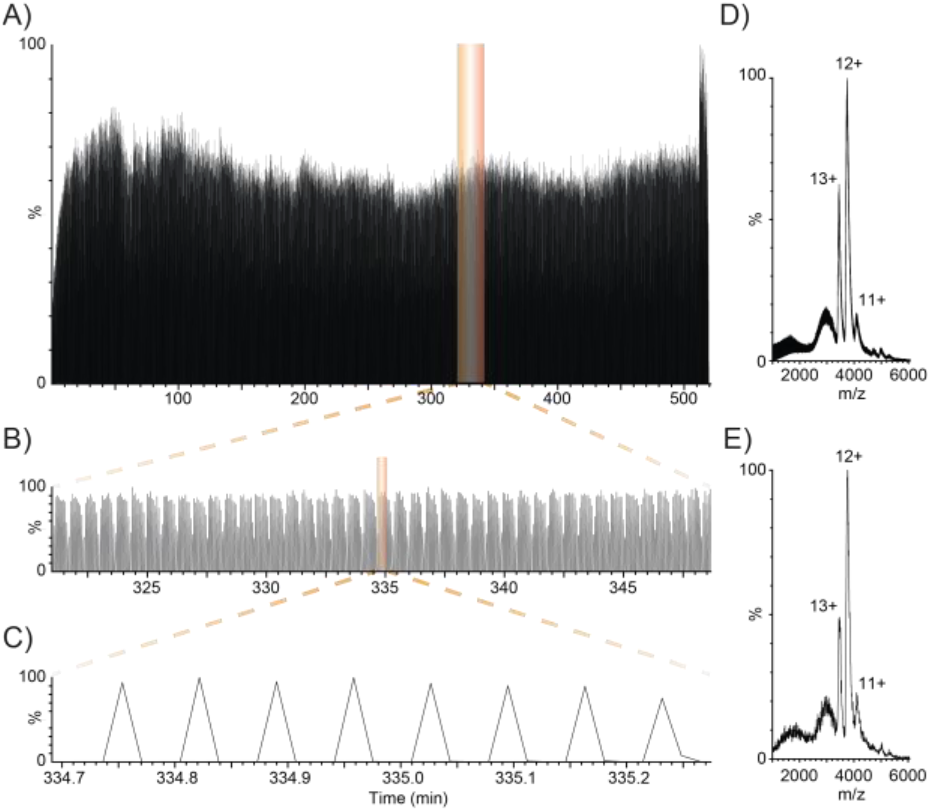
A nESI emitter was loaded with 3 μL of OVA and allowed to be completely consumed while being ionized using TENG. (A) The total time to completely consume the sample was 519 minutes and totaling ∼7,300 pulses results in an average consumption of 414 pL/pulse. TENG provides (B) reproducible peak character and (C) an average of 1 peak every 5 seconds. (D) OVA mass spectra integrated across the entire record length (3 μL) corresponding to the entire 3.50 × 10^−5^ moles of sample being consumed. (E) The mass spectra of OVA integrated across 8 peaks (3.3 nL) produces comparable information content, but only consumed. 3.30 × 10^−14^ moles of sample.

Evaluating the information content more in depth, we do observe a systematic drop off in signal (∼35-50 % from TIC) after ∼7 pulses and believe this may be a mistiming of our TENG device with the ion acquisition time in the trap cell for the proceeding IM experiment (Figure 4B). With optimization we believe that this systematic effect can be minimized or completely negated, and when we zoom into the data further to evaluate the individual TENG peaks outside of this observation, we find that they have great fidelity peak-to-peak in terms of variance of intensity and peak width (Figure 4C). We compared the mass spectrum from OVA averaged using all TENG pulses from the experiment (Figure 4D) and just using just 8 TENG pulses (Figure 4E). While the mass spectrum averaged from just 8 pulses has a lower S/N ratio compared to the full record length of the experiment, overall, both mass spectra have the same information content. When put in context the mass spectrum averaged across 8 peaks would be the equivalent of 3.3 nL of total sample and compute to a total consumption of 3.3 × 10^−14^ moles, or femtomole, quantities of OVA nearly ten orders of magnitude less from the full record length experiment (3.0 × 10^−5^ moles consumed in standard experiment), three orders of magnitude less volume (3 μL in standard experiment) and three orders of magnitude more sensitive than the average quantity of sample that could be obtained from a real cultural heritage sample (nano-picomole range).

To further evaluate the sensitivity of TENG devices, we performed a serial dilution of OVA from the standard 10 μM experiment down to 100 nM (Figure 5A), finding that the characteristic charge states of OVA (11 -13^+^) was retained down to 500 nM without losing fidelity. Below this concentration, significant signal intensity was lost affecting the overall confidence of both qualitative and quantitative evaluations. We hypothesize that this reduction in signal was related to the lower concentrations producing a lower density population of ions from ESI, combined with the high-pressure gases in the trap region of the instrument further reducing the density of the ion packet through scattering collisions. By reducing the gas flow into the trap region (Figure S4, Table S3) we were able to recover the major charge states associated with OVA 11 and 12^+^ charge states but observed significant signal in the 1000 – 3500 m/z range indicating that charge suppression may also play a significant role in the loss of signal at lower concentrations. Additionally, while these low gas pressures may improve signal at low concentrations, they can be detrimental to experiments that require activation in the trap region such as collision induced unfolding most likely due to the gas density being too low to properly confine the ions (Figure S6). Inspired by the recent publication on sub-micron needles from Jordan S.J., *et al*.^28^, we likewise pulled new nESI emitters at 1 pm and 279 nm and confirmed the size of our standard 5 micron emitter and the submicron emitter using SEM (Figure 5B-C, Figure S5). Using these new emitters we were able to simultaneously improve the mass spectrum quality for 100 nM OVA significantly compared to our 5 pm emitters and dramatically reduce the amount of salt clusters present. Lastly, using the new submicron emitter, we obtained CCS measurements for the entire series of concentrations and quantitated that the experimental CCS values obtained are within good agreement (within 3% by error bar) of one another (Figure 4D), and in good agreement (±3%) with the theoretical CCS measurement (Figure 4E). Together, with the previous section on sample volume size and the evaluation of low concentrations required for TENG experiments combined with the structural characterizing capability of IM-MS could provide outstanding analytical measurements for ultra-precious materials such as proteins and protein complexes in objects of cultural heritage.

**Figure 5.**
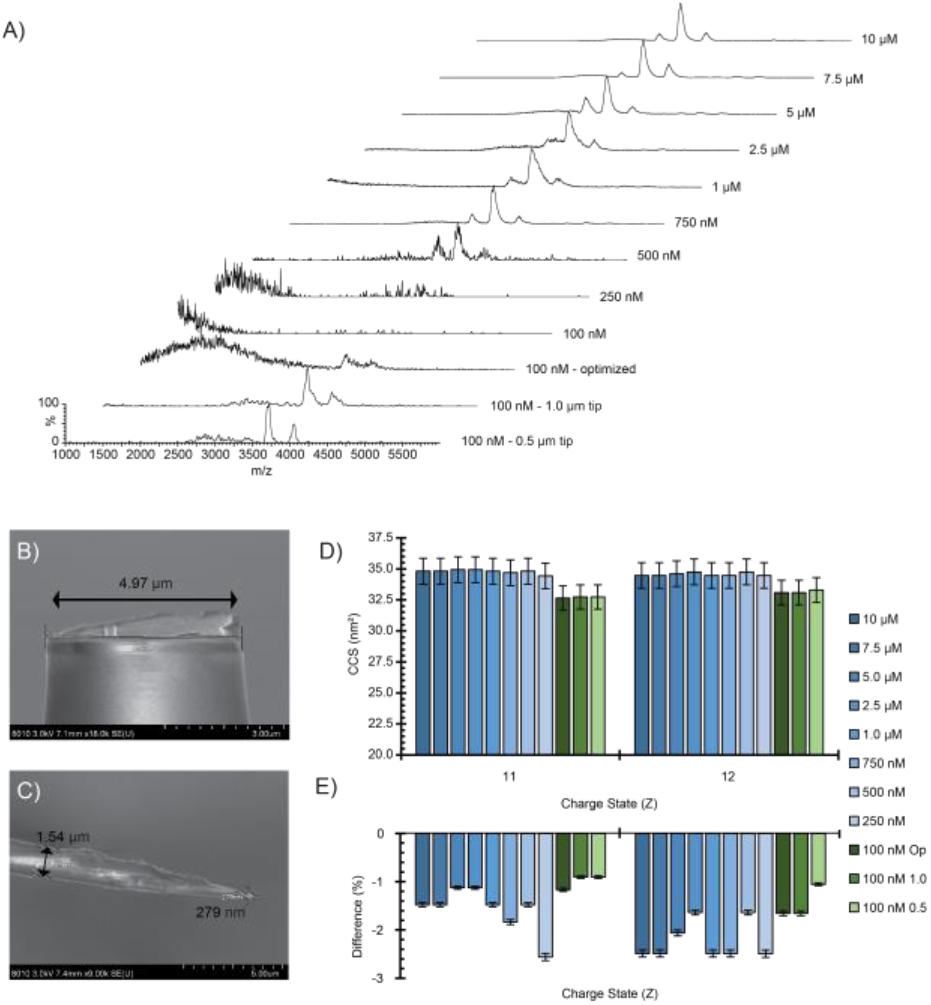
(A) Mass spectral overlay for ovalbumin across a range of concentrations: 10 μM, 7.5 μM, 5 μM, 2.5 μM, 1 μM, 750 nM, 500 nM, 250 nM, and 100 nM. At 250 and 100 nM S/N sharply decreases primarily due to ion suppression from salt clusters. 100 nM signal can be partly recovered through instrument parameter and quadrupole tuning, but salt clusters remain present. The 100 nM signal can be further improved using submicron emitters. SEM images of (B) 5 μm and (C) 0.28 nm emitters. (D) Calculated CCS values for ovalbumin concentration gradient. (E) Percent difference of experimental *vs*. theoretical CCS values, which all fell within 3% for ovalbumin.

## Conclusions

Here, we demonstrate TENG and IM-MS on a series of standard proteins and proteins from eggs with the potential future application for nMS analysis of proteins in cultural heritage objects. To ensure no systematic deviations were observed, three standard proteins were analyzed using DC-based and TENG-based power supplies, which were both calibrated to the respective ion source and cross-calibrated. We found that all three sets of CCS values measured were well within 3% of each other and their corresponding CCS database value. An additional eight standard proteins were used to validate the capabilities of TENG to generate ions under native conditions for a mass range of 5 − 800 kDa, with experimental CCS values being within 3.5% of the database values. Empirically derived theoretical CCS values for a series of egg proteins were compared with our experimental CCS measurements under nMS conditions and a ≤ 3% difference was seen for all except LYS, which likely undergoes a gas-phase compaction due to its lack of HOS. Our findings demonstrate that TENG-based ionization is capable of generating native-like ions, and importantly enables us to access its powerful analytical capabilities as described in previous reports for nMS.

We leveraged the analytical capabilities of TENG by performing a set of serial dilutions and consumption experiments and evaluated those measurements to our results at typical nMS concentrations and volumes. This series of experiments demonstrated that TENG enables volume consumption in the hundreds of pL range, and that signal averaging across only eight TENG peaks, which equates to just 3.3 nL total, resulted in comparable information content in the mass spectrum to the full 3 μL experiment and consumed ten-orders-of-magnitude less sample than traditional nMS experiments. Additionally, we found that for OVA, the concentration could be reduced to 500 nM without significantly effecting the information content of the native mass spectra, and that we could recover the fidelity of samples as low as 100 nM by using sub-micron emitters for the nESI process. These findings combined with previous reports demonstrate that the analytical capabilities of TENG are readily applicable to proteins along with a wide range of molecular classes^13–16^. Moving forward, we aim to apply our TENG-IM-MS technology to model systems mimicking ultra-precious CH samples, which are often only available in the low microgram range and require picomole sensitivity for the proteinaceous fraction. After which, before considering samples from CH samples from paintings and other CH objects will be considered for future study. These results demonstrate that nMS methodologies are suitable to the restrictions of irreplaceable objects and enables state-of-the-art technologies like TENG and IM-MS to enter the CH research toolbox, lifting the lid on the unique and complex structures of proteins within art-matrices and enabling for the first time for the probing of protein degradation mechanisms to advance preservation strategies.

## Supporting information

Supplemental Information Doc

## ASSOCIATED CONTENT

### Supporting Information

The Supporting Information is available free of charge on the ACS Publications website. Detailed information on TENG hardware modifications, comparison of data generated between TENG and DC power supplies, direct infusion consumption experiments, comparison of mass spectra at different gas flows, optical and SEM images of standard and submicron emitters, tables of empirical CCS values compared to experimental measurements, P-97 capillary puller settings, and additional instrumental parameters.

## Author Contributions

Conceptualization, D.D.V., A.P., J.A., C.T., and F.M.F.; Methodology D.D.V and F.M.F.; Formal analysis, D.D.V.; Investigation, D.D.V.; Resources, F.M.F.; Data curation, D.D.V.; Writing—original draft preparation, D.D.V., and A.P.; Writing—review and editing, D.D.V., A.P., J.A., C.T., and F.M.F.; Visualization, D.D.V; Supervision, J.A., C.T., and F.M.F.; Funding acquisition, F.M.F. and D.D.V.; All authors have read and agreed to the published version of the manuscript.

## ACKNOWLEDGMENT

This work is supported by the National Science Foundation Mathematical and Physical Sciences divisions ASCEND program under grant award number CHE-2138107. We would like to thank Bo Yang and the staff at the electrophysiology core as well as the staff at the SEM core for their technical expertise for the fabrication and measurement of the submicron emitters, respectively. We would also like to thank the international laboratory Art and Cultural Heritage (ARCHE) members and the Art Bio Matters community for their collegial support and extensive feedback for the project. Any opinions, findings, and conclusions or recommendations expressed in this material are those of the author(s) and do not necessarily reflect the views of the National Science Foundation.

## Insert Table of Contents artwork here

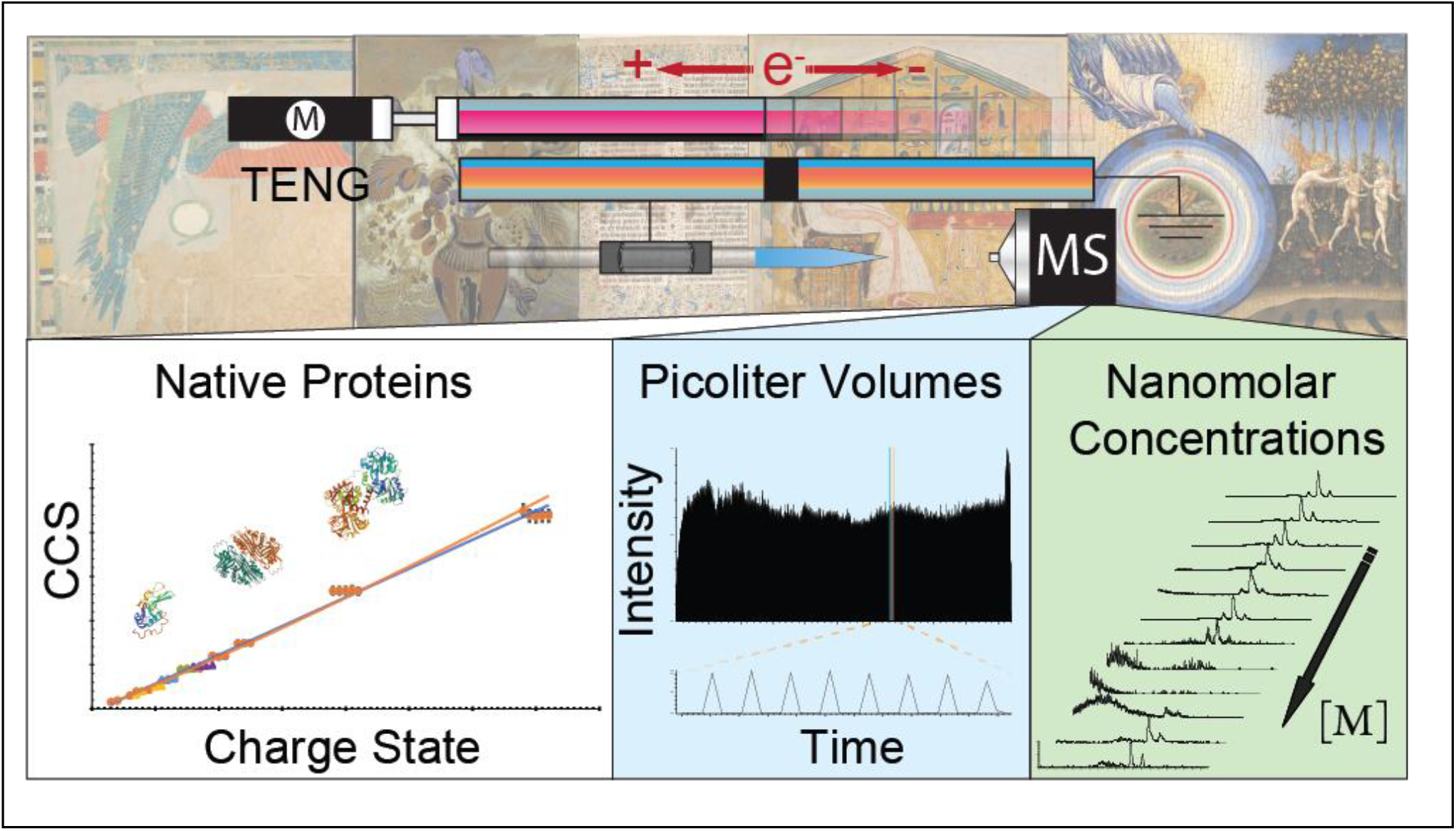

## REFERENCES

(1) Roldán, C.; Murcia-Mascarós, S.; López-Montalvo, E.; Vilanova, C.; Porcar, M. Proteomic and Metagenomic Insights into Prehistoric Spanish Levantine RockArt. Sci. Rep. 2018, 8 (1), 1–10.

(2) Ostwald, W. Letters to a Painter. HW Morse, Boston, New Yor, Chicago, London 1904.

(3) Dallongeville, S.; Garnier, N.; Rolando, C.; Tokarski, C. Proteins in Art, Archaeology, and Paleontology: From Detection to Identification. Chem. Rev. 2016, 116(1), 2–79.

(4) Casadio, F.; Toniolo, L. The Analysis of Polychrome Works of Art: 40 Years of Infrared Spectroscopic Investigations. J. Cult. Herit. 2001, 2 (1), 71–78.

(5) Vandenabeele, P.; Edwards, H. G. M.; Moens, L. A Decade of Raman Spectroscopy in Art and Archeology. Chem. Rev. 2007,107 (3), 675686.

(6) Romero-Pastor, J.; Cardell, C.; Manzano, E.; Yebra-Rodríguez, Á.; Navas, N. Assessment of Raman Microscopy Coupled with Principal Component Analysis to Examine Egg Yolk-Pigment Interaction Based on the Protein C-H Stretching Region (3100-2800 Cm -1). J. Raman Spectrosc. 2011,42 (12), 2137–2142.

(7) Nevin, A.; Osticioli, I.; Anglos, D.; Burnstock, A.; Gather, S.; Castellucci, E. The Analysis of Naturally and Artificially Aged ProteinBased Paint Media Using Raman Spectroscopy Combined with Principal Component Analysis. J. Raman Spectrosc. 2008, 39 (8), 993–1000.

(8) Nevin, A.; Gather, S.; Anglos, D.; Fotakis, C. Analysis of Protein-Based Binding Media Found in Paintings Using Laser Induced Fluorescence Spectroscopy. Anal. China. Acta 2006, 573-574, 341–346.

(9) Vinciguerra, R; Chiaro, A. De; Pucci, P.; Marino, G.; Birolo, L. Proteomic Strategies for Cultural Heritage : From Bones to Paintings. Microchem.J. 2016,126, 341–348.

(10) Dallongeville, S.; Richter, M.; Schafer, S.; Kühlenthal, M.; Garnier, N.; Rolando, C.; Tokarski, C. Proteomics Applied to the Authentication of Fish Glue: Application to a 17th Century Artwork Sample. Analyst 2013, 138(18), 5357–5364.

(11) Pozzi, F.; Arslanoglu, J.; Galluzzi, F.; Tokarski, C.; Snyder, R. Mixing, Dipping, and Fixing: The Experimental Drawing Techniques of Thomas Gainsborough. Merit. Sci. 2020, 8 (1), 1–15.

(12) Galluzzi, F.; Arslanoglu, J.; Tokarski, C. Hydrogen - Deuterium Exchange Mass Spectrometry to Study Interactions and Conformational Changes of Proteins in Paints. Biophys. Chem. 2022, 289.

(13) Li, A.; Zi, Y.; Guo, H.; Wang, Z. L.; Fernández, F. M. Triboelectric Nanogenerators for Sensitive Nano-Coulomb Molecular Mass Spectrometry. Nat. Nanotechnol. 2017, 12 (5), 481–487.

(14) Bernier, M. C.; Li, A.; Winalski, L.; Zi, Y.; Li, Y.; Caillet, C.; Newton, P.; Wang, Z. L.; Fernández, F. M. Triboelectric Nanogenerator (TENG) Mass Spectrometry of Falsified Antimalarials. Rapid Commun. Mass Spectrom. 2018, 32 (18), 1585–1590.

(15) Li, Y.; Bouza, M.; Wu, C.; Guo, H.; Huang, D.; Doron, G.; Temenoff, J. S.; Stecenko, A. A.; Wang, Z. L.; Fernández, F. M. Sub-Nanoliter Metabolomics via Mass Spectrometry to Characterize Volume-Limited Samples. Nat. Commun. 2020, 11 (1), 1–16.

(16) Bouza, M.; Li, Y.; Wu, C.; Guo, H.; Wang, Z. L.; Fernández, F. M. Large-Area Triboelectric Nanogenerator Mass Spectrometry: Expanded Coverage, Double-Bond Pinpointing, and Supercharging. J. Am. Soc. Mass Spectrom. 2020, 31 (3), 727–734.

(17) Ben-Nissan, G.; Sharon, M. The Application of Ion-Mobility Mass Spectrometry for Structure/Function Investigation of Protein Complexes. Curr. Opin. Chem. Biol. 2018, 42, 25–33.

(18) Zhong, Y.; Hyung, S.; Ruotolo, B. T. Ion Mobility-Mass Spectrometry for Structural Proteomics. Expert Rev. Proteomics 2012, 9 (1).

(19) Bush, M. F.; Hall, Z.; Giles, K.; Hoyes, J.; Robinson, C. V; Ruotolo, B. T. Collision Cross Sections of Proteins and Their Complexes: A Calibration Framework and Database for Gas-Phase Structural Biology. Anal. Chem. 2010, 82 (22), 9557–9565.

(20) Hogan, C. J.; Ruotolo, B. T.; Robinson, C. V.; Fernandez De La Mora, J. Tandem Differential Mobility Analysis-Mass Spectrometry Reveals Partial Gas-Phase Collapse of the GroEL Complex. J. Phys. Chem. B 2011,115(13), 3614–3621.

(21) Richardson, K.; Langridge, D.; Dixit, S. M.; Ruotolo, B. T. An Improved Calibration Approach for Traveling Wave Ion Mobility Spectrometry: Robust, High-Precision Collision Cross Sections. Anal. Chem. 2021, 93 (7), 3542–3550.

(22) Haynes, S. E.; Polasky, D. A.; Dixit, S. M.; Majmudar, J. D.; Neeson, K.; Ruotolo, B. T.; Martin, B. R. Variable-Velocity Traveling-Wave Ion Mobility Separation Enhancing Peak Capacity for Data-independent Acquisition Proteomics. Anal. Chem. 2017, 89 (11), 5669–5672.

(23) Polasky, D. A.; Dixit, S. M.; Fantin, S. M.; Ruotolo, B. T. ClUSuite 2: Next-Generation Software for the Analysis of Gas-Phase Protein Unfolding Data. Anal. Chem. 2019, 91 (4), 3147–3155.

(24) Technologies, E. A Simple and Reliable Methodology to Detect Egg White in Art Samples. 2013, 38 (June), 397–408.

(25) Potenza, M.; Sabatino, G.; Giambi, F.; Rosi, L.; Papini, A. M. Analysis of Egg-Based Model Wall Paintings by Use of an Innovative Combined Dot-ELISA and UPLC-Based Approach. 2013, 691–701.

(26) Ujma, J.; Jhingree, J.; Norgate, E.; Upton, R; Wang, X.; Benoit, F.; Beilina, B.; Barran, P. Protein Unfolding in Freeze Frames: Intermediate States Are Revealed by Variable-Temperature Ion Mobility - Mass Spectrometry. 2022.

(27) Valentine, S. J.; Anderson, J. G.; Ellington, A. D.; Clemmer, D. E. Disulfide-Intact and -Reduced Lysozyme in the Gas Phase : Conformations and Pathways of Folding and Unfolding. 1997, 5647 (97), 3891–3900.

(28) Jordan, J. S.; Xia, Z.; Williams, E. R. Tips on Making Tiny Tips: Secrets to Submicron Nanoelectrospray Emitters. J. Am. Soc. Mass Spectrom. 2022, 33 (3), 607–611.

