## Supplemental Information Doc for "Native Triboelectric Nanogenerator Ion Mobility-Mass Spectrometry of Egg Proteins Relevant to Objects of Cultural Heritage at Picoliter and Nanomolar Quantities"

**Table of Contents**

1. Additional information on modifications to TENG device
2. Figure S1: Comparison of ADH TIC, mass spectra, and drift time plots
3. Figure S2: Empirical correlations from database of charge/MW vs. CCS
4. Figure S3: Direct infusion TIC record lengths for consumption experiments
5. Figure S4: Comparison of 100 nM OVA mass spectra at different trap gas flows
6. Figure S5: Optical and SEM images of standard and submicron emitters
7. Figure S6: CIU Fingerprints of OVA
8. Table 1: Empirical CCS values vs experimental CCS measurements
9. Table 2: Sutter P-97 parameters for emitters
10. Table 3: Additional instrument settings

**Modifications to TENG Device**

Updates to the inductive TENG device (TENGi) included the replacement of copper and PTFE with foam and vinyl for the "top" electrode layer, and the replacement of a copper nESI emitter sleeve with a conductive carbon sleeve (Figure 1A). The replacement of the top layer of the sliding electrode with a more pliable material maximized the surface area contact between the top and bottom layers, improving efficiency of triboelectric charge generation and reproducibility of TENG pulses over very long-time scales. Additionally, the original TENGi copper sleeves used to make a connection with the nESI emitter were hand rolled in-house from an adhesive-backed commercial copper sheet. However, it was difficult to achieve consistency between various emitters with these sheets, and the adhesive layer could also interfere in the inductive charging process. The switch to commercial carbon-based conductive foam sleeves (Waters), addressed both of these issues, lowering the entry barrier for TENG device fabrication and operation in other laboratories.

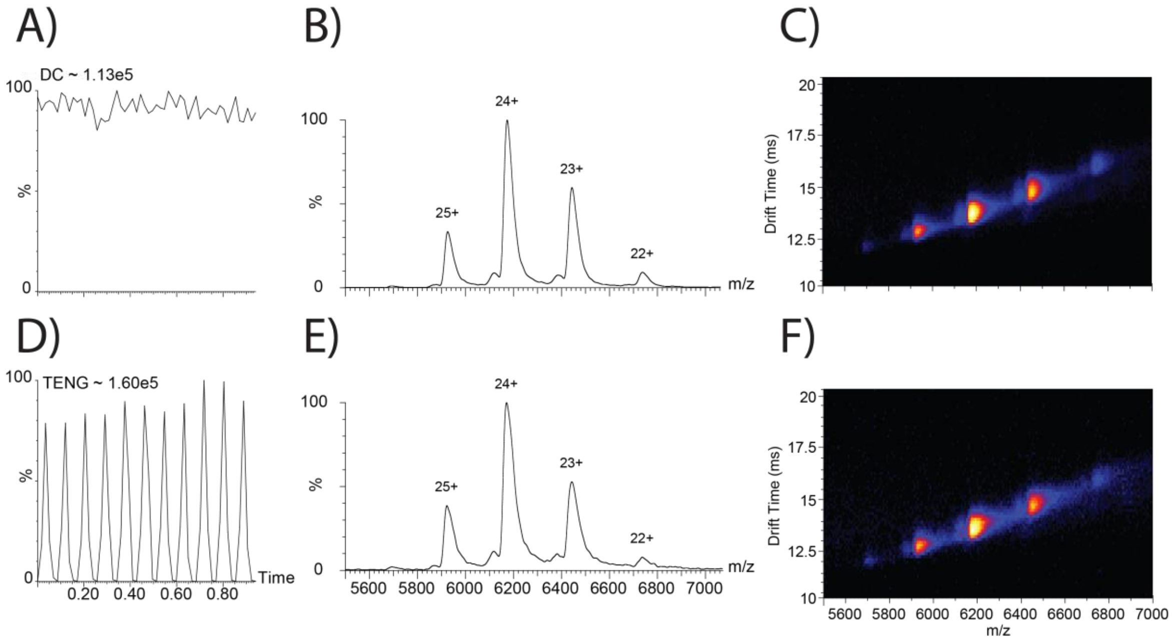


**Figure S1**. Comparison of DC (A) and TENG (D) total ion chronograms, (B,E) mass spectra, and (C,F) drift time plots for alcohol dehydrogenase, showing comparable information content.

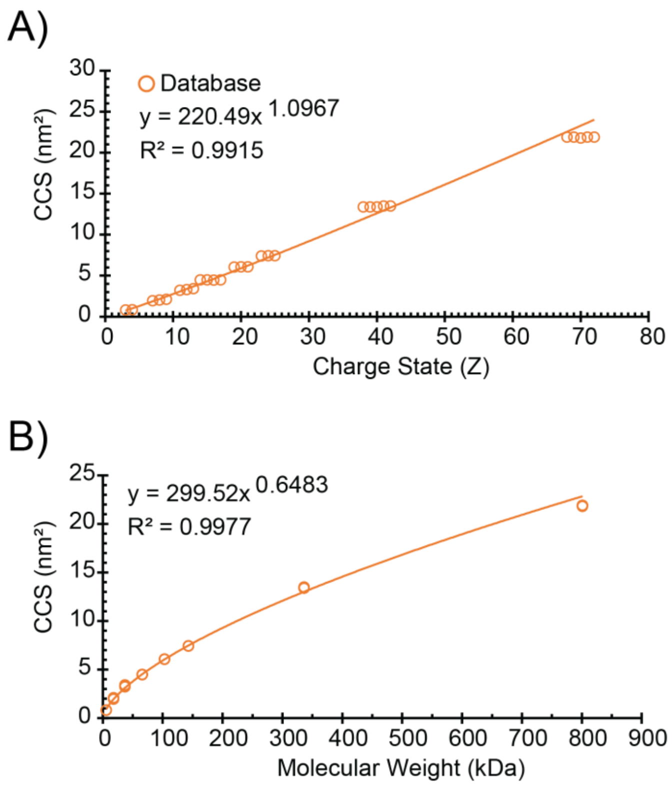


**Figure S2**. Empirical correlation plots of database CCS vs. (A) charge state and (B) molecular weight in kDa.

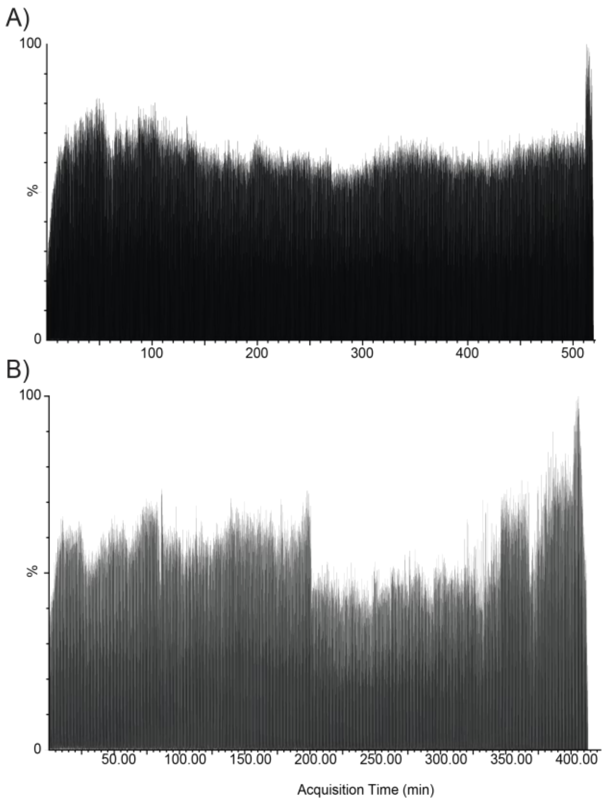


**Figure S3**. Volume consumption experiments using (A) 3 µL and (B) 2 µL of ovalbumin sample.

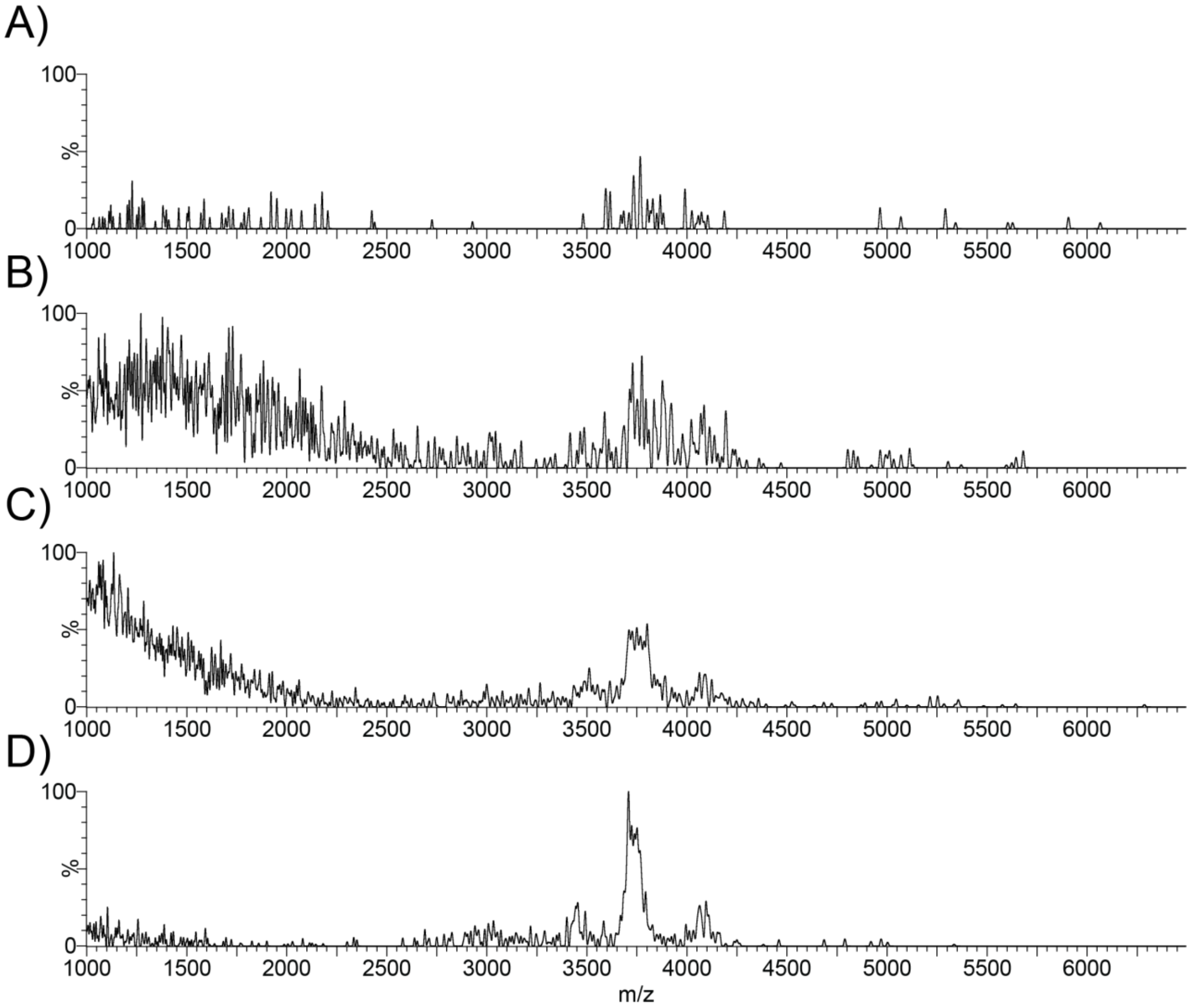


**Figure S4**. Comparison of OVA at 100 nM concentrations at trap gas flow rates of (A) 10, (B) 5, (C) 3, and (D) 1 (mL/min) .

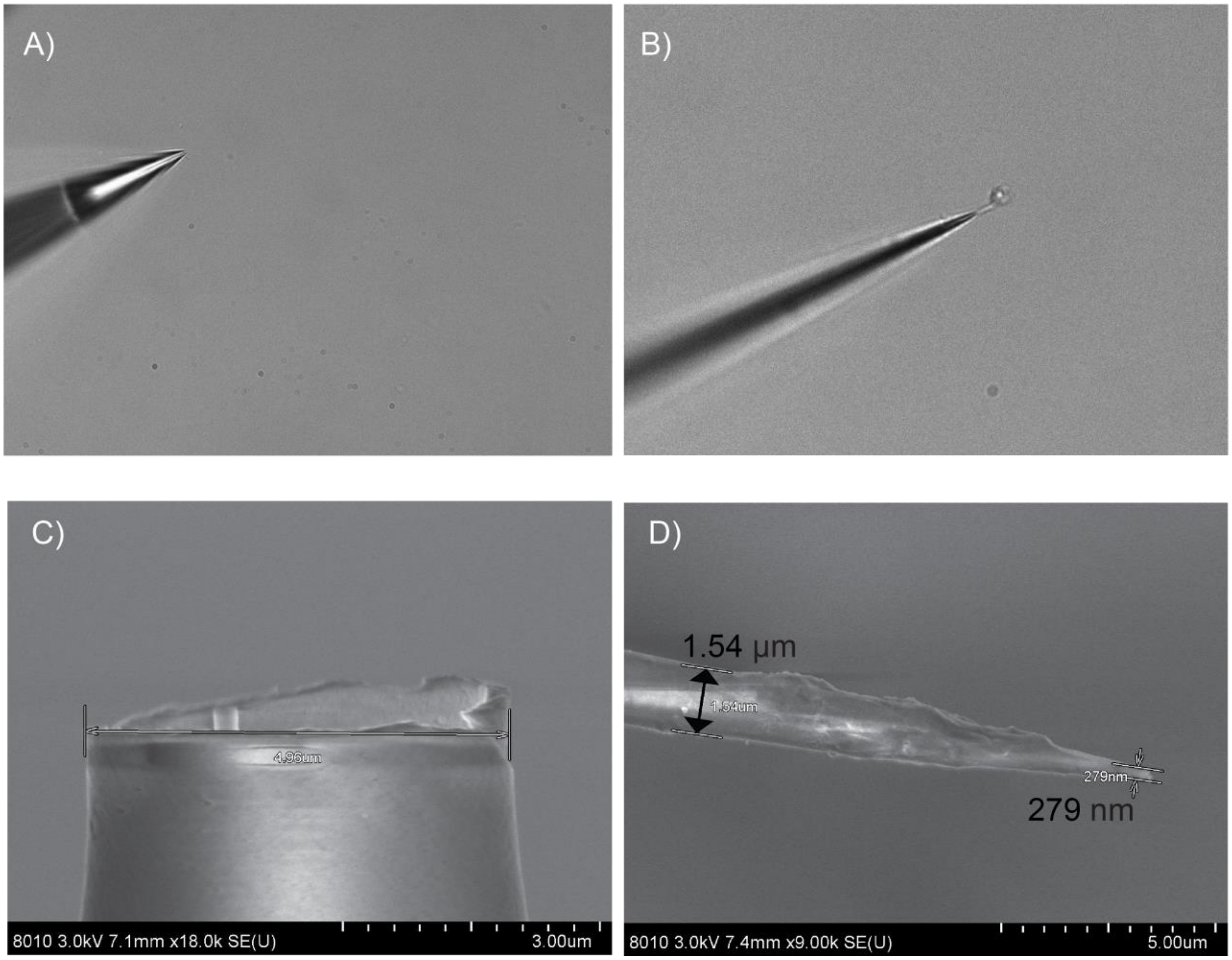


**Figure S5**. Comparison of 5 (A,C) and .279 (B,D) µm emitters using 40x objective microscope (A,B) and scanning electron microscopy (C,D).

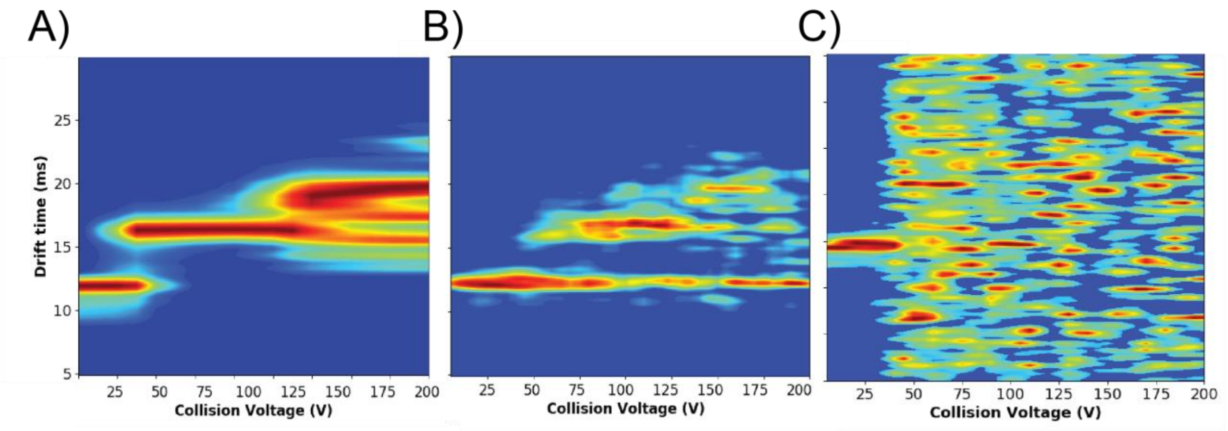


**Figure S6**. Comparison of CIU fingerprints for OVA at (A) 10 µM, (B) 100 nM and trap gas flow of 3, and (C) 100 nM and trap gas flow of 1 (mL/min).

**Table 1.** CCS measurements for standard proteins compared to CCS database and CCS values calculated from molecular weight and charge empirically derived from correlations from CCS database values. Green highlighted cells indicate the CCS value of standard proteins used for % difference analysis.

| Standard Protein | m / kDa | Z | $\Omega_{N2}$ (nm$^2$) | DB $\Omega_{N2}$ (A$^2$) | Z Calculated $\Omega_{N2}$ (A$^2$) | $M_w$ Calculated $\Omega_{N2}$ (A$^2$) | Average | Std. Dev. | % Difference DB | % Difference Z | % Difference MW |
|---|---|---|---|---|---|---|---|---|---|---|---|
| Insulin | 5.8 | 3 | 8.2513 | 825.1 | 735.6 | 936.2 | 763.60 | 18.13 | -7.75 | 3.73 | -20.30 |
| Insulin* | 5.8 | 4 | 8.492 | 849.2 | 1008.5 | 936.2 | 923.80 | 9.16 | 8.41 | -8.77 | -1.33 |
| CytC | 12 | 6 | 14.9 | 1490.0 | 1573.2 | 1499.9 | 1484.94 | 7.89 | -0.34 | -5.77 | -1.00 |
| CytC | 12 | 7 | 15.9 | 1590.0 | 1863.0 | 1499.9 | 1620.40 | 29.80 | 1.89 | -13.93 | 7.72 |
| β-lac | 18 | 7 | 19.5 | 1950.0 | 1863.0 | 1950.8 | 1934.30 | 46.29 | -0.81 | 3.76 | -0.85 |
| β-lac | 18 | 8 | 20.3 | 2030.0 | 2156.8 | 1950.8 | 1990.14 | 9.58 | -1.98 | -8.04 | 2.00 |
| β-lac | 18 | 9 | 21.1 | 2110.0 | 2454.2 | 1950.8 | 2186.49 | 15.99 | 3.56 | -11.54 | 11.39 |
| β-lac | 37 | 11 | 32.3 | 3230.0 | 3058.3 | 3112.4 | 3090.59 | 16.69 | -4.41 | 1.05 | -0.70 |
| β-lac | 37 | 12 | 33.1 | 3310.0 | 3364.5 | 3112.4 | 3227.58 | 46.90 | -2.52 | -4.16 | 3.63 |
| β-lac | 37 | 13 | 34.3 | 3430.0 | 3673.3 | 3112.4 | 3369.50 | 39.83 | -1.78 | -8.63 | 7.93 |
| HSA | 66 | 14 | 44.9 | 4490.0 | 3984.3 | 4529.4 | 4407.79 | 72.45 | -1.85 | 10.09 | -2.72 |
| HSA | 66 | 15 | 44.9 | 4490.0 | 4297.4 | 4529.4 | 4474.67 | 87.31 | -0.34 | 4.04 | -1.21 |
| HSA | 66 | 16 | 44.7 | 4470.0 | 4612.6 | 4529.4 | 4541.95 | 85.27 | 1.60 | -1.54 | 0.28 |
| HSA | 66 | 17 | 44.9 | 4490.0 | 4929.7 | 4529.4 | 4614.82 | 91.86 | 2.74 | -6.60 | 1.87 |
| ConA | 103 | 19 | 60.6 | 6060.0 | 5569.3 | 6044.3 | 5867.54 | 57.99 | -3.23 | 5.22 | -2.97 |
| ConA | 103 | 20 | 60.8 | 6080.0 | 5891.5 | 6044.3 | 5899.52 | 58.81 | -3.01 | 0.14 | -2.42 |
| ConA | 103 | 21 | 60.9 | 6090.0 | 6215.4 | 6044.3 | 5946.58 | 42.97 | -2.38 | -4.42 | -1.63 |
| ADH | 143 | 23 | 74.2 | 7420.0 | 6867.5 | 7477.0 | 7409.61 | 97.11 | -0.14 | 7.59 | -0.91 |
| ADH | 143 | 24 | 74.5 | 7450.0 | 7195.6 | 7477.0 | 7428.43 | 81.65 | -0.29 | 3.18 | -0.65 |
| ADH | 143 | 25 | 74.4 | 7440.0 | 7525.1 | 7477.0 | 7490.45 | 81.06 | 0.68 | -0.46 | 0.18 |
| GDH | 336 | 38 | 134 | 13400.0 | 11910.7 | 13009.2 | 13338.13 | 50.72 | -0.46 | 11.31 | 2.50 |
| GDH | 336 | 39 | 134 | 13400.0 | 12254.9 | 13009.2 | 13326.23 | 75.60 | -0.55 | 8.38 | 2.41 |
| GDH | 336 | 40 | 134 | 13400.0 | 12599.9 | 13009.2 | 13351.50 | 4.60 | -0.36 | 5.79 | 2.60 |
| GDH | 336 | 41 | 135 | 13500.0 | 12945.8 | 13009.2 | 13354.78 | 80.09 | -1.08 | 3.11 | 2.62 |
| GDH | 336 | 42 | 135 | 13500.0 | 13292.5 | 13009.2 | 13414.33 | 72.62 | -0.64 | 0.91 | 3.07 |
| GroEL | 801 | 68 | 219 | 21900.0 | 22547.7 | 22848.1 | 22381.13 | 97.49 | 2.17 | -0.74 | -2.07 |
| GroEL | 801 | 69 | 219 | 21900.0 | 22911.6 | 22848.1 | 22166.70 | 101.26 | 1.21 | -3.30 | -3.03 |
| GroEL | 801 | 70 | 218 | 21800.0 | 23276.0 | 22848.1 | 22058.40 | 0.00 | 1.18 | -5.37 | -3.52 |
| GroEL | 801 | 71 | 219 | 21900.0 | 23640.9 | 22848.1 | 22068.70 | 107.48 | 0.77 | -6.88 | -3.47 |
| GroEL | 801 | 72 | 219 | 21900.0 | 24006.3 | 22848.1 | 22052.77 | 220.71 | 0.70 | -8.48 | -3.54 |

**Table 2.** Parameters used for a Sutter Instruments P-97 Flaming Brown Puller for preparing submicron emitters with diameters of approximately 5, 1, and 0.28 µm.

| Diameter (µm) | Ramp Temp | Heat | Pull | Velocity | Delay | Cycles |
|---|---|---|---|---|---|---|
| 5 | 524 | 582 | - | 30 | 1 | 1 |
| 1 | 524 | 592 | - | 26 | 1 | 1 |
| 0.28 | 524 | 533 | 70 | 80 | 150 | 1 |

**Table 3.** Synapt G2 settings used for all experiments. Highlighted parameters associated with the "optimized" settings for 100 nM experiments with standard (e.g. 5 μm emitters). Values obtained from _extern.inf files.

| Parameter | Standard | "Optimized" |
|---|---|---|
| **Source Temperature** | 30 | 30 |
| **Sampling Cone** | 40 | 40 |
| **Extraction Cone** | 0 | 0 |
| **Trap Collision Energy** | 10 | 10 |
| **Trap Gas Flow** | 3-6 | 1 |
| **IMS Wave Velocity** | 600 | 600 |
| **IMS Wave Height** | 40 | 40 |
| **Backing Pressure** | 3.21e0 | 3.07e0 |
| **Source Pressure** | 2.35e-3 | 2.37e-3 |
| **Trap Pressure** | 3.10e-2 | 2.45e-2 |
| **IMS Pressure** | 4.06e0 | 3.98e0 |
| **Transfer Pressure** | 1.00e-6 | 1.00e-6 |
| **TOF Pressure** | 1.35e-6 | 1.12e-6 |
| **Acquisition Range** | | |
| **Start mass** | 1000 | 1000 |
| **End mass** | 12000 | 12000 |
| **Acquisition Time (mins)** | 2 | 2 |
